# Analysis of site and structure specific core fucosylation in liver disease progression using exoglycosidase-assisted data-independent LC-MS/MS

**DOI:** 10.1101/2020.07.29.227488

**Authors:** Miloslav Sanda, Jaeil Ahn, Petr Kozlik, Radoslav Goldman

**Affiliations:** Department of Oncology, Lombardi Comprehensive Cancer Center, Georgetown University, Washington, D.C. 20057; Department of Biochemistry and Molecular & Cellular Biology, Georgetown University, Washington, D.C., 20057; Department of Biostatistics, Bioinformatics & Biomathematics, Georgetown University Medical Center, Washington, D.C., 20057; Clinical and Translational Glycoscience Research Center, Georgetown University, Washington, D.C., 20057; Department of Analytical Chemistry, Faculty of Science, Charles University, Prague, Czech Republic

**Keywords:** Mass spectrometry, Data independent acquisition, GP-SWATH, Core fucosylation, Soft fragmentation, N-glycopeptide, Cirrhosis

## Abstract

Carbohydrates form one of the major groups of biological macromolecules in living organisms. Many biological processes including protein folding, stability, immune response, and receptor activation are regulated by glycosylation. Fucosylation of proteins regulates such processes and is associated with various diseases including autoimmunity and cancer. Mass spectrometry efficiently identifies structures of fucosylated glycans or sites of core fucosylated N-glycopeptides but quantification of the glycopeptides remains less explored. We performed experiments that facilitate quantitative analysis of the core fucosylation of proteins with partial structural resolution of the glycans and we present results of the mass spectrometric SWATH-type DIA analysis of relative abundances of the core fucosylated glycoforms of 45 glycopeptides derived from 18 serum proteins in liver disease of different etiologies. Our results show that a combination of soft fragmentation with exoglycosidases is efficient at the assignment and quantification of the core fucosylated N-glycoforms at specific sites of protein attachment. In addition, our results show that disease-associated changes in core fucosylation are peptide-dependent and further differ by branching of the core fucosylated glycans. Further studies are needed to verify whether tri- and tetra-antennary core fucosylated glycopeptides could be used as markers of liver disease progression.

## INTRODUCTION

N-glycosylation is a frequent posttranslational modification of proteins with significant influence on physiological and pathological processes ^1-5^. Fucose (deoxygalactose) can be attached directly to serines (S) and threonines (T) of some proteins ^6^ and is a common building block of glycans in the N- and O-glycoproteins ^7^. Two distinct positional isomers are described in the context of fucosylated human N-glycoproteins. One set of fucosylated isomers, described as outer-arm or terminal fucosylation, is attached to the antennae of the complex type N-glycans by α-(1-3) and α-(1-4) linkage to the GlcNAc residues, by α-(1-2) linkage to galactose, or by α-(1-4) linkage to GalNAc ^8^. The second positional isomer called core fucose is attached by α-(1-6) linkage to the GlcNAc bound to the protein’s asparagine ^6^. This positional isomer is formed in glycoproteins by the transfer of fucose from GDP-fucose to the innermost GlcNAc residue exclusively by the α-(1-6)-fucosyltransferase (Fut8) ^9,10^.

Core fucosylation is widespread and all classes of human N-glycans (paucimannose, high mannose, hybrid, complex) are core fucosylated to some extent ^11-13^. It is also an impactful modification. Studies of Fut8(-/-) knockout mice ^14^ or congenital disorders of glycosylation ^15^ show that lack of this essential modification leads to severe health consequences. Mechanisms of the pathophysiological processes are often unknown as core fucosylation affects many molecular pathways. Perhaps best defined function of the core fucose is modified affinity of the glycosylated IgG for FcγRIII ^16^ which affects immune responses or the function of therapeutic antibodies ^17-19^. A number of additional functions of core fucosylation was described in the studies of growth factor signaling ^14^ or T-cell receptors ^20,21^ but there is much to be discovered.

Alterations in these pathways were associated with various types of cancer ^22-25^, including hepatocellular carcinoma ^23^, as well as other diseases ^26,27^. It is therefore not surprising that a number of mass spectrometric methods were introduced to study the extent of core fucosylation and its association with the disease processes ^28-33^. In this study, we introduce a combination of exoglycosidase digests with data independent (DIA) LC-MS/MS using low collision energy (CE) fragmentation to achieve quantification of the core fucosylated glycopeptides with partially resolved N-glycan structures. We demonstrate applicability of the method to biolomedical problems on the relative quantification of the changes in core fucosylation of 45 glycopeptides derived from 18 serum proteins of patients with liver cirrhosis of various etiologies.

## EXPERIMENTAL SECTION

### Materials and Reagents

Acetonitrile (LC-MS grade), water (LC-MS grade), and dithiothreitol (DTT) were obtained from ThermoFisher Scientific (Waltham, MA). Iodoacetamide (IAA) was from MP Biomedicals (Santa Ana, CA). Trypsin Gold (V5280) was obtained from Promega (Madison, WI). α2-3,6,8,9 neuraminidase (P0722), α1-2 (P0724) and α1-3,4 (P0769) fucosidases with GlycoBuffer 1 (5 mM calcium chloride, 50 mM sodium acetate, pH 5.5) were purchased from New England BioLabs (Ipswich, MA). Hemopexin from human plasma was supplied by Athens Research and Technology (Athens, Georgia). Rapigest SF was from Waters (Milford, MA). All other chemicals were obtained from SigmaAldrich (Saint Louis, MO).

### Study Population

Applicability of the method was documented on serum samples of healthy volunteers and patients with cirrhosis of the liver of hepatitis C (HCV), hepatitis B (HBV), alcoholic (ALD) and non-alcoholic steatohepatitis (NASH) etiologies as described previously ^34^. Briefly, participants were enrolled in collaboration with the Department of Hepatology and Liver Transplantation, Georgetown University Hospital, Washington, DC, under protocols approved by the Institutional Review Board. Blood samples were collected using red-top serum vacutainer tubes (BD Diagnostics, Franklin Lakes, NJ) in line with manufacturer’s recommendations and samples were processed within 6 h of the blood draw for storage at -80 °C. We created two pooled samples for each disease category. Each pool represents five participants and we analyzed duplicate samples of each of the pools (4 samples total per group). All the pools were age-matched and had a comparable degree of liver damage, as measured by the model for end-stage liver disease (MELD) score.

### Sample Processing

Affinity depletion of abundant serum proteins was carried out on a HP1290 HPLC system (Agilent, Santa Clara, CA) using the Multiple Affinity Removal Column Human 14 (MARS 14, Agilent) according to manufacturer’s protocol. Briefly, human serum (20 μl) was mixed with buffer A (85 μl), remaining particulates were removed by centrifugation through a 0.22-μm spin filter 1 min at 16,000 × g, and the clarified serum was injected on the column. Affinity buffer A and B were used as mobile phases, 100% A at a flow rate 0.25 ml/min at 0-9 min, 100% B at a flow rate 1 ml/min at 9-15 min, and 100% A at a flow rate 1 ml/min at 15-20 min. Collected flow-thru of each sample was precipitated using 80% acetone overnight at -20 °C, washed 2-times with 100% acetone, and dissolved in 50 mM ammonium bicarbonate buffer pH 8 for determination of the protein concentration by BCA assay. Aliquots of the samples (10 μg of total protein) were reduced with 5 mM DTT for 60 min at 60°C, alkylated with 15 mM IAA for 30 min in the dark, residual IAA was quenched with 5 mM DTT, and digested with 2.5 ng/μl Trypsin Gold (Promega, Madison, WI) at 37°C in Barocycler NEP2320 (Pressure BioSciences, South Easton, MA) for 1 hour. Trypsin was deactivated by heating to 99 °C for 10 minutes, samples were evaporated using a vacuum concentrator (Labconco, Kansas City, MO) and digested with 2 μL of α2-3,6,8,9 neuraminidase in GlycoBuffer 1 overnight at 37 °C according to manufacturer’s instructions. Neuraminidase was deactivated by heating at 75 °C for 10 minutes and 2 μL aliquots of each 1-3,4 fucosidase and 1-2 fucosidase were added to the samples for an overnight digestion at 37 °C. The digests were evaporated using a vacuum concentrator (Labconco) and reconstituted at a final concentration 0.5 µg/µL in a mobile phase A (0.1% formic acid in 2% acetonitrile) for mass spectrometric analysis.

### Glycopeptide identification by nano LC-MS/MS

Following 5 min trapping/washing step in solvent A (2% acetonitrile, 0.1% formic acid) at 10 μL/min, peptide and glycopeptide separation was achieved at a flow rate of 0.3 μL/min using the following gradient: 0-3 min 2% B (0.1% formic acid in ACN), 3-5 min 2-10% B; 5-60 min 10-45% B; 60-65 min 45-98% B; 65-70 min 98% B, 70-90 min equilibration at 2% B. We used an Orbitrap Fusion Lumos mass spectrometer to analyze the glycopeptides with the electrospray ionization voltage at 3 kV and the capillary temperature at 275 °C. MS1 scans were performed over m/z 400–1800 with the wide quadrupole isolation on a resolution of 120,000 (m/z 200), RF Lens at 40%, intensity threshold for MS2 set to 2.0e4, selected precursors for MS2 with charge state 3-8, and dynamic exclusion 30s. Data-dependent HCD tandem mass spectra were collected with a resolution of 15,000 in the Orbitrap with fixed first mass 110 and normalized collision energy 10-30%. LC-MS datasets were processed by Protein Discoverer 2.2. (Thermo Fisher Scientific, Waltham, MA) with Byonic node (Protein Metrics, Cupertino, CA) followed by manual confirmation of the glycopeptides.

### SWATH DIA quantification by nano LC-MS/MS

Glycopeptide separation was achieved on a Nanoacquity LC (Waters, Milford, MA) using capillary trap, 180 μm x 0.5 mm, and analytical 75 μm x 150 Atlantis DB C18, 3 μm, 300 Å columns (Water, Milford, MA) interfaced with 6600 TripleTOF (Sciex, Framingham, MA). A 3 min trapping step using 2% ACN, 0.1% formic acid at 15 µl/min was followed by chromatographic separation at 0.4 µl/min as follows: starting conditions 5% ACN, 0.1% formic acid; 1-55 min, 5–50% ACN, 0.1% formic acid; 55-60 min, 50–95% ACN, 0.1% formic acid; 60-70 min 95% ACN, 0.1% formic acid followed by equilibration to starting conditions for additional 20 min. For all runs, we have injected 1 µl of tryptic digest on column. We have used Data Independent Acquisition (DIA) with one MS1 full scan (400-2000 m/z) and n MS/MS fragmentations (800-1600), dependent on the isolation window (15 Da), with rolling collision energy-optimized as described previously ^35^. MS/MS spectra were recorded in the range 400-2000 *m/z* with resolution 30,000 and mass accuracy less than 15 ppm using the following experimental parameters: declustering potential 80V, curtain gas 30, ion spray voltage 2,300 V, ion source gas1 11, interface heater 180°C, entrance potential 10 V, collision exit potential 11 V. Precursor and product ion masses and charge states together with the glycopeptide retention times (RT) used for quantification of the glycoforms are summarized in **Supplemental Table 1**.

### Data analysis

Y-ion isotope clusters with isolation window of 1.2 Da, extracted from the low CE SWATH-MS/MS with a 15 Da step window, were used for analysis of the glycopeptide intensities. Multiquan 2.0 software was used for processing of the quantitative results. Processing methods were created for ion extraction from each SWATH window and for each glycoform. Areas of the fucosylated glycopeptides were normalized to the areas of the corresponding non-fucosylated analytes. Coefficients of variability (CV) were determined from the means and standard deviations of the measurements and were compared across the disease groups and glycoform categories using one-way analysis of variance (ANOVA). Normalized glycopeptide intensities (measured as peak areas) and their ratio’s were compared in the different disease groups by Kruskal-Wallis one-way ANOVA with statistical significance determined as a two-sided p<0.05. Statistical analyses were performed using R version 3.40; further data processing and graphing were carried out in Microsoft Excel and Graphpad prism.

## RESULTS

### Glycosidase treatment and identification of the N-glycopeptides in human serum

Major goal of our study was to evaluate the feasibility of DIA quantification of core fucosylated glycopeptides (GP-SWATH) without an endoglycosidase digest. We do this to evaluate whether changes in the core fucosylation of liver secreted glycoproteins observed in fibrotic liver disease differ with branching of the glycoforms. To this end, we analyzed tryptic digests of MARS14 depleted human serum. The glycopeptides were further simplified by sequential treatment with exoglycosidases (α2-3,6,8,9 neuraminidase followed by α1-2 and α1-3,4 fucosidases) and were subsequently analyzed by LC-MS/MS without any further glycopeptide enrichment. We verified completeness of the exoglycosidase digests by nano HILIC separation of the hemopexin glycopeptide **(Supplemental Figure 1)** as described previously ^36^. This assures that we quantify core fucosylated glycoforms of each peptide but allows comparison of fucosylation on glycoforms with different degree of branching (see below).

We prepared pooled samples of healthy controls and patients with cirrhosis of the liver caused by HCV, HBV, ALD, and NASH etiologies. For each group, we prepared two pooled samples (n=5 participants each) which we analyzed first by LC-MS/MS with HCD fragmentation on an Orbitrap Fusion Lumos as described above. We selected for further analysis bi-, tri-, and tetra-antennary glycoforms of 45 N-glycopeptides of 18 proteins detectable in all the samples in their fucosylated and non-fucosylated forms **(Supplemental Table 1)**. In addition to the 24 bi-antennary glycoform pairs (HexNAc4Hex5Fuc/HexNAc4Hex5), our analysis included 13 glycopeptides with tri-antennary (HexNAc5Hex6Fuc/HexNAc5Hex6) glycoforms, and ten glycopeptides with tetra-antennary (HexNAc6Hex7Fuc/HexNAc6Hex7) glycoforms. In total, we quantified 90 glycopeptides by the DIA LC-MS/MS workflow.

### Core fucosylation and limited fragmentation

To facilitate the DIA analysis of glycopeptides, we used lower CE settings (half of the CE used for peptides) to minimize interference of the peptides, to minimize rearrangement of fucose and unwanted fragmentation of the glycans, and to maximize the yield of the quantification fragments as described previously ^35,37^. Transitions for the quantification of the 90 glycopeptides use the Y-ions corresponding, typically, to the loss of an outer-arm of the glycan; this generates a Y-ion of [m/z-1] compared to the precursor and the corresponding oxonium ion. The low CE fragmentation of the core fucosylated and non-fucosylated glycopeptide pairs is quite similar as documented on the spectra of the bi-antennary glycopeptide SWPAVGNCSSALR of hemopexin **(Figure 1)**. The spectra of both the fucosylated **(Figure 1a)** and non-fucosylated **(Figure 1b)** glycoforms contains Y-ions (m/z 1404.6 fucosylated and m/z 1331.6 non-fucosylated) that represent the loss of an arm (singly charged HexNAc-Hex m/z 366.1) and Y-ions (m/z 1323.6 and m/z 1250.5) from the loss of one arm with manose (singly charged HexNAc-Hex-Hex m/z 528.2) visible in the low mass end. The m/z 366.1 oxonium ions are not seen because we acquire the spectra from 400 Da due to optimized ion transmission for the Y-ions.

**Figure 1.**
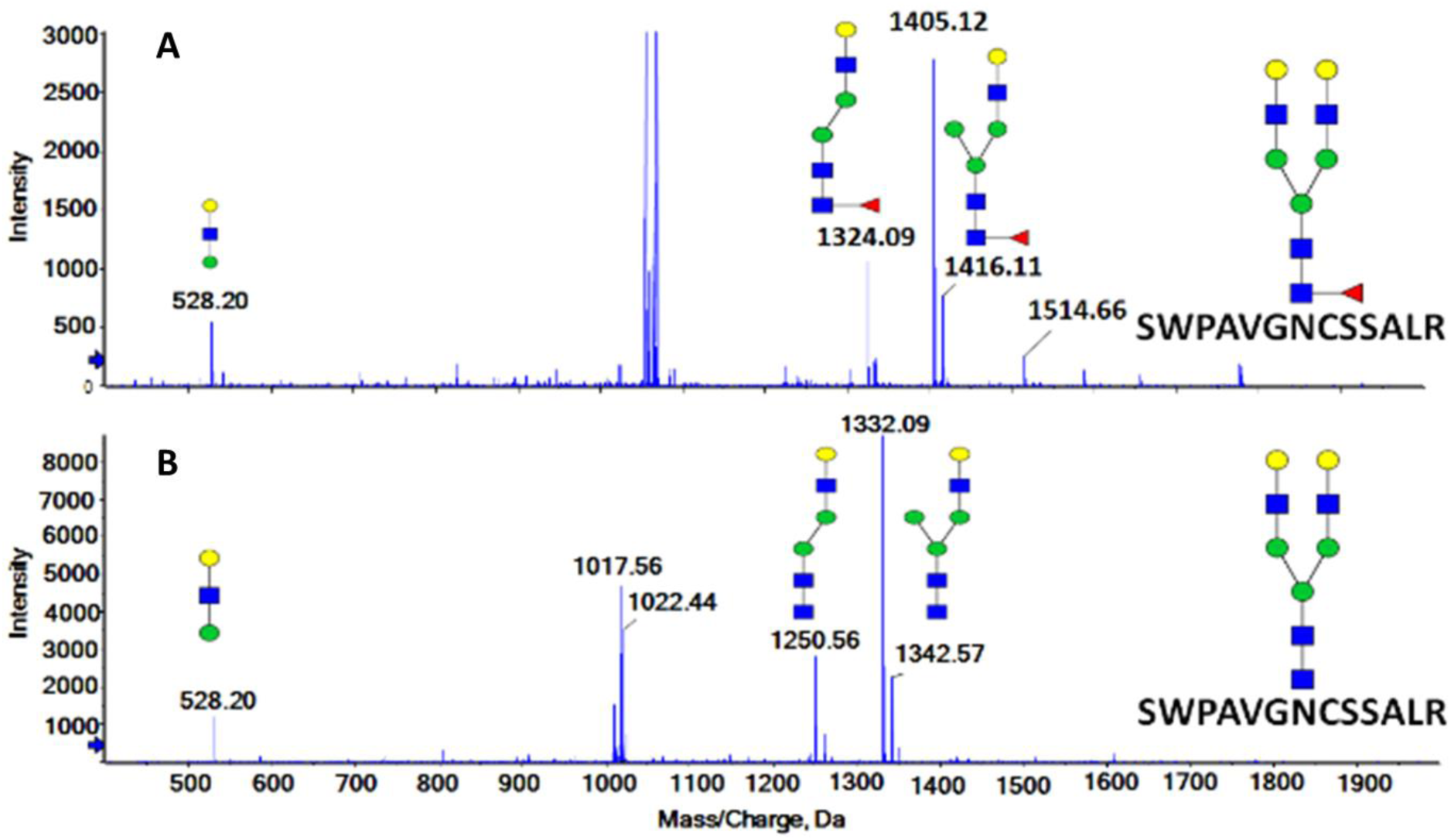
Low collision energy spectra from the GP-SWATH DIA of the following glycoforms of the SWPAVGNCSSALR glycopeptides of hemopexin: (A) bi-antennary core fucosylated glycoform; and (B) bi-antennary glycoform without fucose.

The low CE fragmentation allows efficient extraction of the product ions for quantification of the glycoforms as documented on the SWPAVGNCSSALR glycopeptide of hemopexin **(Figure 2)**. The XIC chromatograms show the expected trend of slight decrease in the retention time (RT) of the glycoforms with the addition of each neutral carbohydrate unit as described previously ^38^. The RT alignment further confirms the identity of the quantified analytes. The low CE settings also minimize the potential rearrangement of fucose which keeps the error in quantification of the fucosylated glycoform below 15% ^37,39^ and maximizes the ion intensity sufficiently for reliable quantification of even the low-abundant tetra-antennary glycoforms.

**Figure 2.**
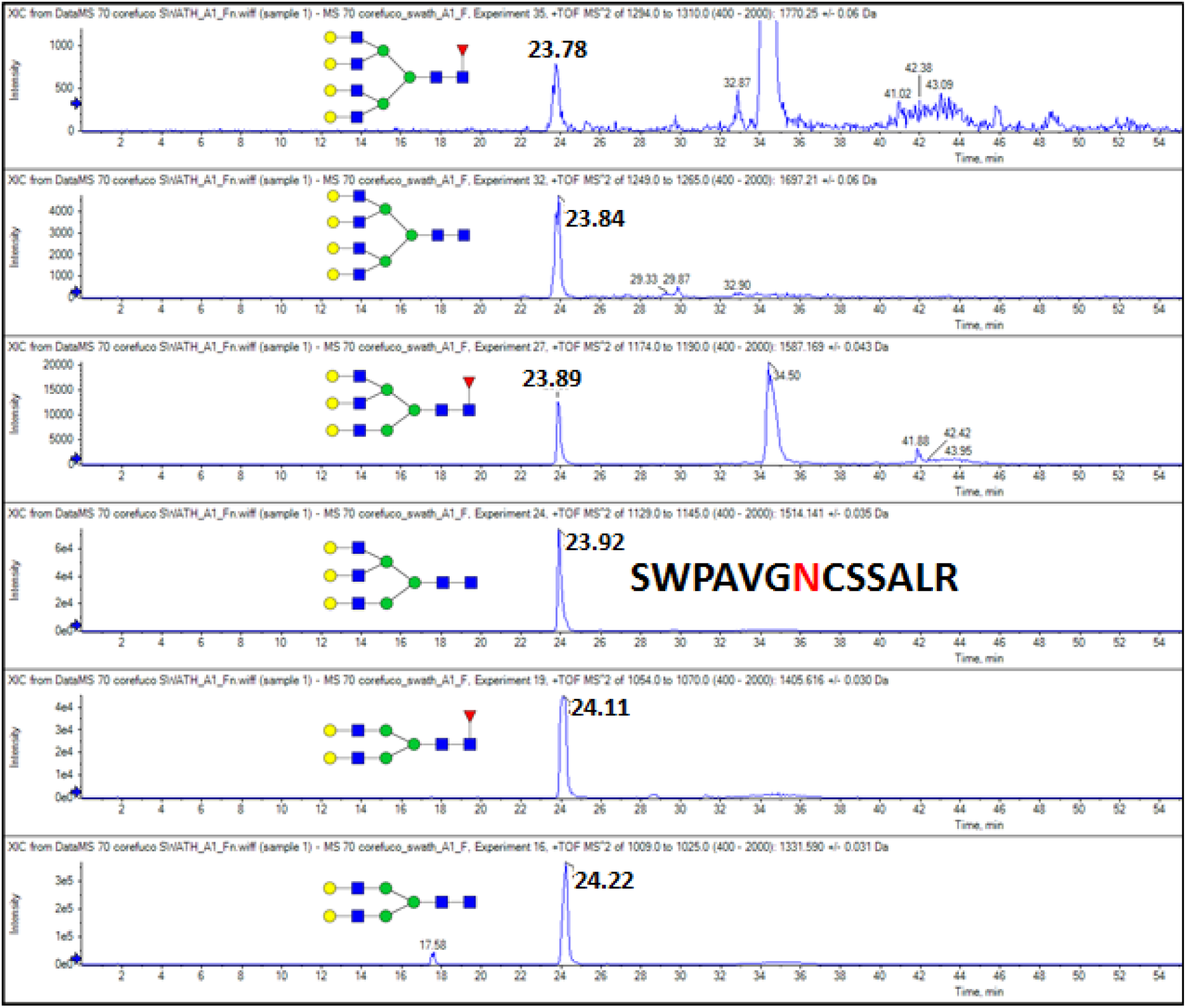
XIC chromatogram of six glycoforms of the SWPAVGNCSSALR glycopeptide of hemopexin extracted from the GP-SWATH DIA data of a MARS14 depleted serum sample treated with α2-3,6,8,9 neuraminidase, α1-2 and α1-3,4 fucosidases.

This is documented by the evaluation of the coefficients of variation (CV) across all the 90 glycopeptides quantified; we summarize the results as average CVs in the respective disease groups and glycoform classes **(Table 1)**. We measured every sample in duplicate and we compare the duplicate measurements to derive the technical CVs **(Table 1A)**. The average technical CVs of all fucosylated or non-fucosylated glycopeptides are 16.9% and 18.2%, respectively, and they are not statistically different (one-way ANOVA p=0.925). Our analysis shows that the technical CVs do not differ between the disease groups but the CVs of tetra-antennary nonfucosylated glycoforms are higher compared to the tri- or bi-antennary analytes. Interestingly technical CVs of fucosylated tetraantennary glycoforms does not follow this trend even if they are more abundant than nonfucosylated and influence of noise could be lower. As expected, the biological replicates (two different samples per group) show similar CVs with average CV 23.3% for the fucosylated glycopeptides compared to the average CV of 22.0% for the non-fucosylated glycopeptides (one-way ANOVA p=0.216). We do not observe a significant differences in the biological CVs by disease group (one-way ANOVA p=0.091) or by the number of <u>arms (</u>one-way ANOVA p=0.445). Higher biological variability, observed especially in the disease groups, is expected and is minor compared to the fold-changes observed between the healthy controls and the cirrhotic disease groups (see below).

**Table 1.** Distribution of the average coefficients of variability (CV) for the indicated glycopeptide classes and disease groups across the entire study set. Duplicate measurements were performed for both the (A) technical and (B) biological CV measures.

|  |  |  |  |  |  |
| --- | --- | --- | --- | --- | --- |
| A. Technical CV |  |  |  |  |  |
| Alcoholic | HBV | HCV | NASH | Control | Glycan |
| 16.39 | 15.92 | 17.95 | 13.01 | 14.24 | FA2 |
| 17.46 | 21.84 | 19.60 | 14.34 | 20.40 | FA3 |
| 16.71 | 17.93 | 17.44 | 14.27 | 16.90 | FA4 |
| 15.23 | 13.95 | 13.27 | 11.75 | 13.97 | A2 |
| 15.14 | 22.74 | 16.06 | 11.35 | 14.96 | A3 |
| 24.00 | 30.16 | 22.55 | 24.85 | 23.33 | A4 |
| B. Biological CV |  |  |  |  |  |
| Alcoholic | HBV | HCV | NASH | Control | Glycan |
| 28.25 | 24.99 | 32.91 | 27.34 | 23.89 | FA2 |
| 17.98 | 27.58 | 31.33 | 13.49 | 13.85 | FA3 |
| 29.00 | 16.37 | 22.66 | 7.37 | 20.09 | FA4 |
| 17.28 | 33.28 | 19.50 | 16.37 | 16.76 | A2 |
| 17.51 | 30.78 | 24.25 | 14.00 | 15.40 | A3 |
| 23.11 | 27.88 | 26.50 | 13.93 | 11.96 | A4 |

### Trends in the abundance of fucosylated and non-fucosylated glycopeptides

The core fucosylated glycopeptides show a general trend to increased intensities in the cirrhotic groups compared to the healthy controls **(Supplemental Table 1)**. When we compare peak areas of the same fucosylated glycoforms in the cirrhosis groups to the healthy controls, 23 of 45 glycopeptides have higher intensity in cirrhosis; 6 of the glycopetides show significantly higher intensity in all four cirrhosis categories (p<0.05) compared to the healthy controls and all of them have intensity consistently more than 2-fold higher. Only 2 core fucosylated glycopeptides show consistently lower intensities in the four cirrhosis groups compared to the healthy controls. The highest difference in the peak areas of the fucosylated glycopeptide was observed for the tetra-antennary glycoform of the ELHHLQEQNVSNAFLDK glycopeptide of ceruloplasmin which is on average 7.5 fold higher in the cirrhosis groups. One of the consistenly and significantly changed core fucosylated glycopeptides is the biantennary and triantennary LANTQGEDQYYLR peptide of clusterin (**Figure 3**) which was previously described in liver disease by our group^33^ and by others (panel of analytes in patent US20090208926A1).

**Figure 3.**
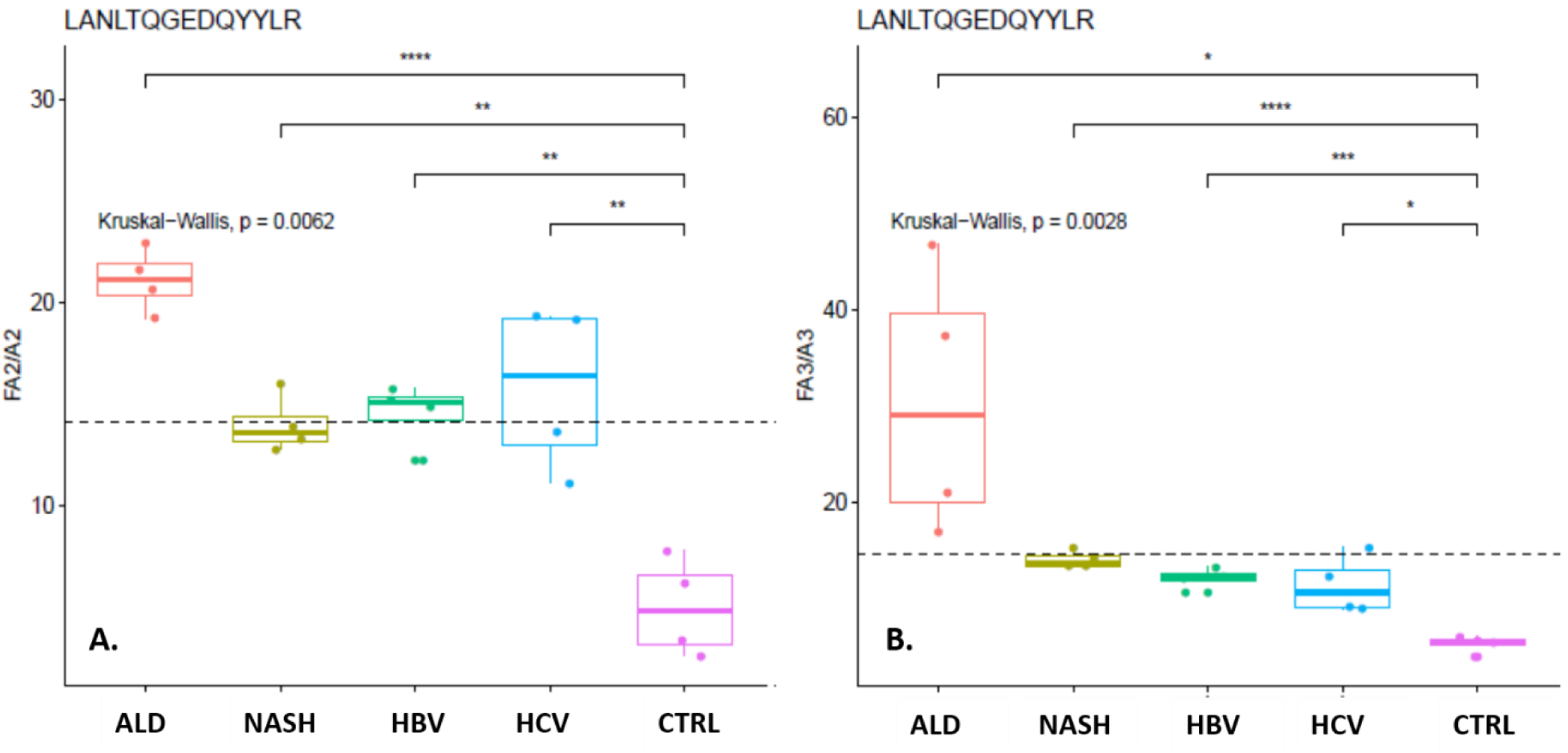
Changes in the relative abundance of the (A) bi-antennary and (B) tri-antennary core fucosylated glycoforms of the LANTQGEDQYYLR peptide of clusterin in healthy controls and patients with liver cirrhosis of the indicated etiologies.

At the same time, we observe a trend to lower intensities of the non-fucosylated glycopeptides in the cirrhosis groups compared to the healthy controls **(Supplemental Table 1)**. We observe a consistently lower intensity of 14 glycopeptides in the four cirrhosis groups (p<0.05) and one of them has intensities decreased more than 2-fold. Only 3 non-fucosylated glycopeptides show significantly higher intensity in at least 2 of the cirrhosis groups. This trend reflects to some degree the reported trends in the serum concentration of the proteins in liver cirrhosis. We observe lower intensities in the non-fucosylated glycopeptides of proteins (hemopexin, clusterin, ITIH4, fibrinogen) reported to decrease in cirrhosis while the non-fucosylated glycopeptide of alpha-2 macroglobulin has higher intensity in cirrhosis in line with its increased serum concentration ^40-42^. However, additional factors affect the abundance of the non-fucosylated glycopeptides because some glycoforms of the same peptide show inconsistent trends. For example, the tri- and tetra-antennary glycoforms of some glycopeptides have higher intensity in cirrhosis while their bi-antennary glycoforms are lower. This trend may be potentially explained by increased branching in the cirrhosis groups. This is likely the case of the ELHHLQEQNVSNAFLDK glycopeptide of ceruloplasmin where the tetra-antennary non-fucosylated glycoform has higher intensity in the cirrhosis groups while the bi-antennary glycoform is slightly lower in cirrhosis than in the healthy controls.

These trends suggest that the abundance of the core fucosylated glycoforms of some proteins goes up in cirrhosis in spite of the fact that the protein’s concentration may decrease. It is also a reason why we normalize the fucosylated glycoforms by their non-fucosylated counterparts and consider ratio as the two glycoforms as our final measure of interest.

### Structure specific changes of core fucosylation in liver cirrhosis

We find multiple core fucosylated glycoproteins with significantly higher intensities of fucosylated glycopeptides in the serum of some category of the cirrhotic disease. Core fucosylated glycoforms (normalized intensity) of 12 glycopeptides of these proteins increase significantly (p<0.05) 3-fold or greater in at least two categories of liver cirrhosis compared to the healthy controls **(Table 2)**. Here we compare the normalized fucosylated intensities represented by the ratios of the same fucosylated/non-fucosylated glycoforms of a glycopeptide (i.e. bi-antennary fucosylated/bi-antennary non-fucosylated etc). Overall, we observe the highest (5- to 9-fold) increase in the normalized fucosylated intensity for the bi-antennary glycoform of the QVFPGLNYCTSGAYSNASSTDSASYYPLTGD glycopeptide of apolipoprotein B-100; the increase is significant in all the cirrhosis groups compared to the healthy control group. However, we also observe decrease in the normalized fucosylated intensities for some glycopeptides. For example, the bi-antennary FGCEIENNR glycopeptide of Zn-alpha2-glycoprotein has significantly lower normalized fucosylated intensities in the cirrhosis groups (Supplemental Table 1) in line with our previous reports of site- and structure-specific changes in the fucosylation of serum proteins in the context of liver cirrhosis ^35,43^. In addition, we observe an interesting trend towards higher intensities of branched core fucosylated glycoforms for several peptides. For example, the fucosylated intensities of the tri-antennary glycoform of the SWPAVGNCSSALR peptide of hemopexin increase in cirrhosis consistently 2- to 5-fold while the intensities of the bi-antennary glycoforms increase 1- to 3-fold. Equally interesting are the trends of the bi-, tri- and tetraantennary glycoforms of the VCQDCPLLAPLNDTR peptide of Alpha-2-HS-glycoprotein and the LGNWAMPSCK glycopeptide of beta-2-glycoprotein **(Figure 4)**. For all three glycopeptides, the increase in the intensity of the fucosylated analyte goes up with branching. We do not know the reason for this observation; it could be related to different rates of synthesis or to the post-synthetic selection of the fucosylated glycoforms. Nonetheless, the results show clearly that the presented DIA LC-MS/MS workflow enables quantification of the partially resolved fucosylated structures which would not be achieved with the commonly used truncation of the glycoforms with endoglycosidases.

**Table 2.**
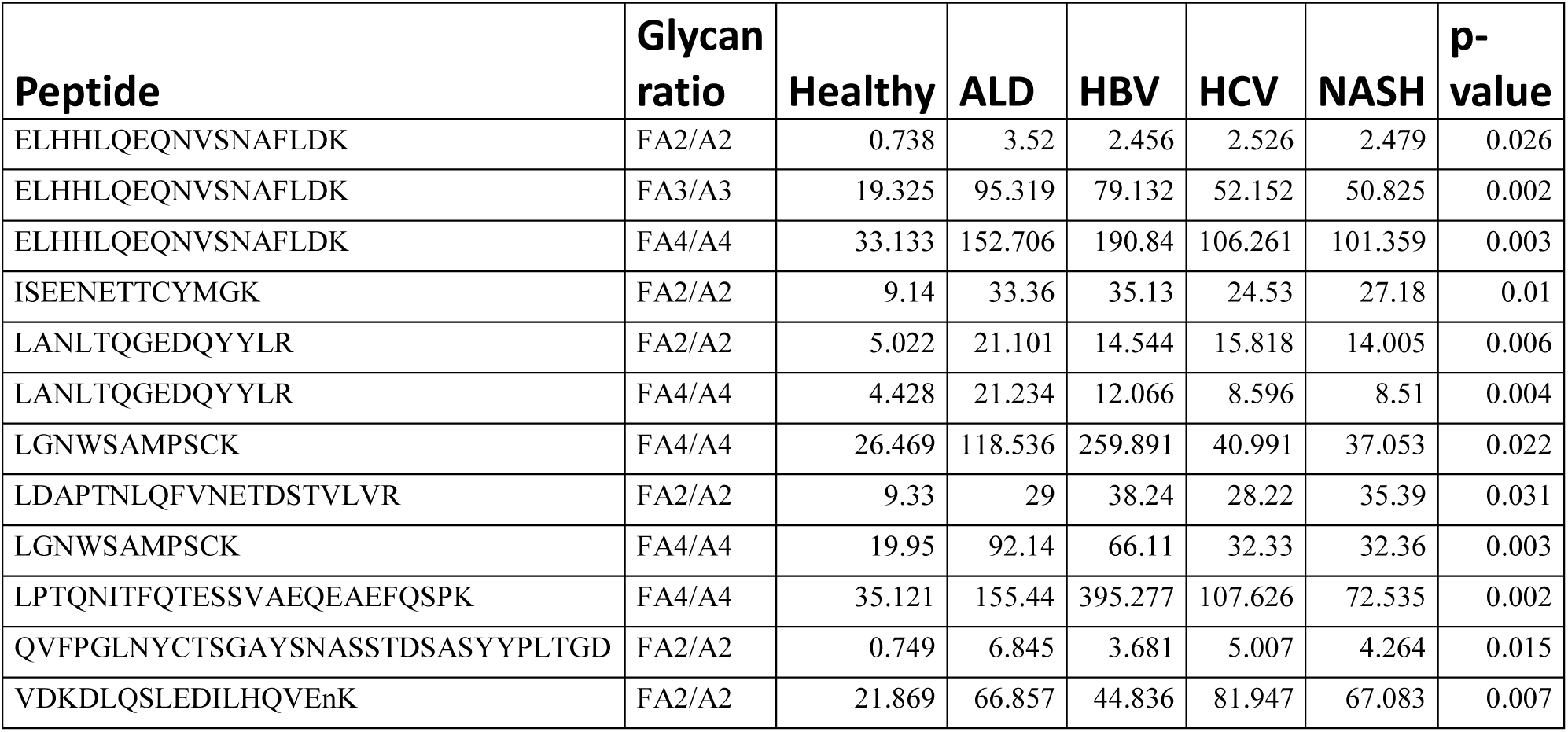
Normalized intensities of the core fucosylated glycoforms of peptides with ratio of the glycoforms increased more than 3-fold in at least two categories of cirrhotic liver disease compared to the healthy controls.

**Figure 4.**
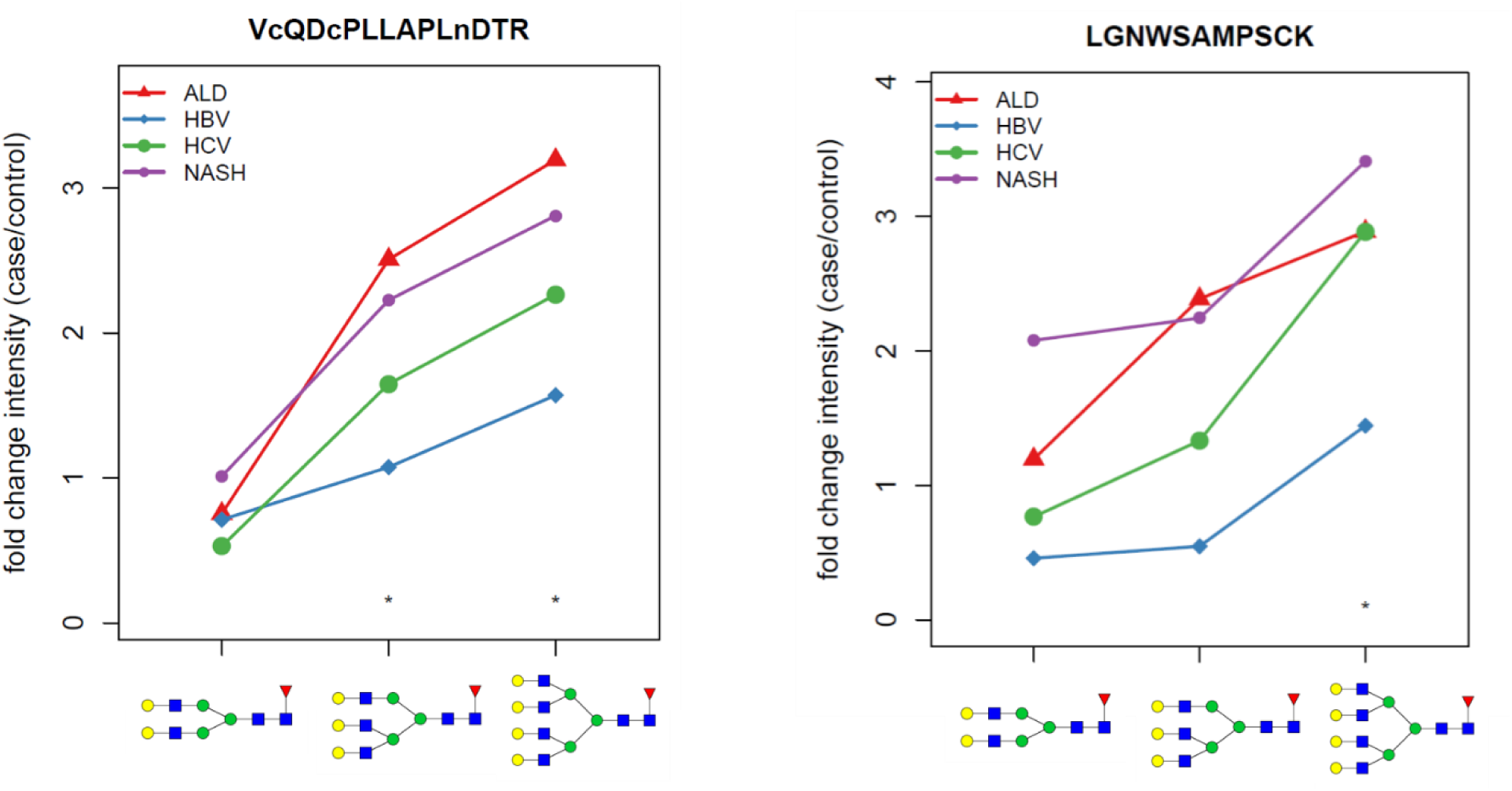
Ratio of the increase in the intensity of the following core fucosylated glycopeptides in the liver cirrhosis groups of four etiologies compared to the healthy controls: A. bi-, tri- and tetra-antennary glycoforms of the VCQDCPLLAPLNDTR peptide of Alpha-2-HS-glycoprotein; B. bi-, tri-, and tetra-antennary glycoforms of the LGNWAMPSCK peptide of beta-2-glycoprotein.

## DISCUSSION

Knowledge of the extent of core fucosylation of human proteins substantially expanded in recent years ^28-33^. The reported studies use a variety of enrichment strategies and mass spectrometric workflows to improve coverage of the site-specific core fucosylated proteins but quantification of the core fucosylated analytes remains challenging ^28-33^. This is due to some extent to the complex nature of quantification of the glycoforms at specific sites of attachment to proteins ^44^ but also due to the fact that synthetic isotopically labeled standards are not available for the quantification of glycopeptides. The studies therefore focus on relative quantification using iTRAQ or TMT labeling ^30-32^ which is often combined with truncation of the N-glycans with endoglycosidases ^29,31^ to improve identification and relative quantification of the truncated (peptide with GlcNAc with/out fucose) analytes. This is, however, by no means straightforward as the efficiency of cleavage of the iTRAQ or TMT tags is affected by the presence of complex glycans and because the rate of cleavage of some types of complex glycans (fucosylated, branched) by endoglycosidases differs substantially (some are not appreciably cleaved). As a consequence, we know of several thousand sites modified to some extent by core fucosylated N-glycans but we know their relative intensities to a limited extent ^28-33^.

In this study, we introduce a workflow that eliminates the endoglycosidase digest and extends the glycopeptide SWATH (GP-SWATH) analyses, that we introduced recently ^35^, to relative quantification of the core fucosylated glycopeptides. We combine exoglycosidase digests with DIA LC-MS/MS using low CE fragmentation which enables quantification of the core fucosylated glycopeptides with partially resolved N-glycan structures, namely branching into bi-, tri-, and tetra-antennary glycoforms. This is important because the knowledge of the core fucose distribution on glycoproteins by their branching is limited.

We use exoglycosidases (α2-3,6,8,9 neuraminidase, 1-3,4 fucosidase and 1-2 fucosidase) because we focus on the core fucose. The outer-arm modifications dilute the MS signal into multiple low abundant analytes which may have overlapping isotope clusters ^45^; removal of the outer-arm fucose also simplifies data interpretation. Low CE fragmentation is an important component of the workflow because it minimizes interferences of peptides (which remain to a large extent unfragmented), eliminates excessive fragmentation of the glycans **(Figure 1)** and potential rearrangement of fucose as described recently ^39,46^. This set of optimizations makes the glycopeptide Y-ions sufficiently selective for reliable quantification of the branched (bi- to tetra-antennary) core fucosylated glycopeptides by the GP-SWATH DIA LC-MS/MS. Our results document relative quantification of 45 glycopeptide pairs (with/out fucose) with average technical CV around 17% **(Table 1)** and demonstrate meaningful biological assessment of differences in the core fucosylation of serum glycoproteins in liver cirrhosis of various etiologies.

We observe typically higher core fucosylation in liver cirrhosis compared to the healty controls; a typical increase is up to 2-fold but we observed up to 8-fold increased intensity of the fucosylated glycoform in case of the tetra-antennary ELHHLQEQNVSNAFLDK glycopeptide of ceruloplasmin, depending on the etiology of the cirrhosis. The increase in core fucosylation is more pronounced in the groups of cirrhosis of ALD (average: 1.79) and NASH (average 1.88) etiologies (15 and 21 of the 45 glycopeptides with significantly higher core fucosylation compared to the healhy controls, respectively) as opposed to HBV and HCV etiology (8 and 12 glycopeptides). At the same time, we observe a trend towards lower intensity of the non-fucosylated glycoforms in cirrhosis which is likely related, to some degree, to the decrease in concentration of some of the proteins in the cirrhotic patients. It is well known that synthesis of proteins in the liver is affected by the progressing fibrotic remodeling. We therefore normalize the intensity of the fucosylated glycoforms and use the ratio of fucosylated to the non-fucosylated forms as the quantified readout **(Table 2)**.

Some bi-antennary glycoppetides decrease in cirrhosis compared to healthy controls but increase in cirrhosis in the tri- or tetra-antennary forms of the same peptide. This cannot be explained by lower protein concentration but could perhaps be attributed to increased branching; further studies will be needed to define whether this originates at synthesis or post-synthetic selection. The VC(cam)QDC(cam)PLLAPLNDTR glycopeptide of Alpha-2-HS-glycoprotein is a good example with higher contribution of tetra-antennary glycoforms in the cirrhosis groups while the bi-antennary and triantennary glycoforms remain the same. Relative contribution of the bi-, tri and tetra-antennary structres in the healthy group for this peptide is 18%,78% and 4%, respectively and theratio of the glycoforms in the cirrhotic groups compared to the heatlhy group range from 0.71 to 1.04 for the bi-antennary glycopeptides, 0.98 to 1.05 (for tri-antennary, and 1.20 to 1.40 for the tetra-antennary glycopeptides. This is true for several glycopeptides (for example LGNWSAMPSCK), and supports the usefulness of our analytical approach where we use increase of core fucosylation and increase of glycan branchinch to simultaneously. The greater than 4-fold increases of the normalized intensities of some of the glycopeptides in cirrhosis suggests that the core fucosylated glycopeptides might be usefull in serologic monitoring of liver fibrosis; this hypothesis is, however, by no means confirmed by our preliminary results and needs to be further evaluated.

Our study has several weaknesses. We rely on the quantification of relative intensities using a label-free DIA workflow because we do not have synthetic isotopic standards of the glycopeptides. Nonetheless, the workflow has an average technical CV of 17.5% and shows consistent trends which suggests good performance. In addition, we do not have yet an automated pipeline for the analysis of the GP-SWATH DIA datasets which will be needed to improve the throughput of the studies; this is an active topic of our ongoing studies.

In summary, we introduce an exoglycosidase-assisted GP-SWATH analysis of core fucosylated glycopeptides with resolved branching of the N-glycans. We show that the DIA LC-MS/MS workflow identifies meaningful differences in the relative quantities of site-specific core fucosylated glycopeptides in a preliminary study of liver cirrhosis. The data suggest that the degree of core fucosylation and its disease-associated alterations differ by branching of the N-glycans as documented on the case of ceruloplasmin or hemopexin. We expect that the workflow will enable quantitative studies of core fucosylation in biological model systems and in liquid biopsies such as serum, urine, or cerebrospinal fluid (CSF).

## Abbreviations

(S): serine
(T): threonine
(CE): collision energy
(DIA): data independent acquisition
(GP-SWATH): glycopeptide SWATH
(HCV): hepatitis C virus
(HBV): hepatitis B virus
(ALD): alcoholic
(NASH): non-alcoholic steatohepatitis.

## Acknowledgments

This work was supported by National Institutes of Health Grants U01CA230692, R01CA135069, and R01CA238455 (to RG); further support was provided by the Office Of The Director, National Institutes of Health under Award Number S10OD023557 supporting the operation of the Clinical and Translational Glycoscience Research Center, Georgetown University, CCSG Grant P30 CA51008 (to Lombardi Comprehensive Cancer Center) supporting the Proteomics and Metabolomics Shared Resource, and the Czech Science Foundation, Grant No 19-18005Y (to PK). The content is solely the responsibility of the authors and does not necessarily represent the official views of the National Institutes of Health.

