## Supplemental Material for "Analysis of site and structure specific core fucosylation in liver disease progression using exoglycosidase-assisted data-independent LC-MS/MS"

**Supplemental table 1:** Summary of DIA results of selected fucosylated glycopeptides in liver disease

| Peptide sequence | Glycan composition (after treatment with exoglycosidases) glycan with/out core fucose | m/z precursor non fucosylated | m/z precursor fucosylated | charge (z) precursor | Quantification fragment non-fucosylated | Quantification fragment fucosylated | RT (min) | Intensity Healthy | Intensity ALD | Intensity HBV |
|---|---|---|---|---|---|---|---|---|---|---|
| | | | | | | | | Fucosylated | | |
| AALAAFNAQNNGSNFQLEEISR | HexNAc4Hex5Fuc/HexNAc4Hex5 | 1329.916 | 1378.602 | 3 | 1811.805 | 1884.834 | 34.49 | 1124213.1 | 1237895 | 957536 |
| AFITNFSMIIDGMTYPGIIK | HexNAc4Hex5Fuc/HexNAc4Hex5 | 1285.58 | 1334.266 | 3 | 1745.3 | 1818.329 | 34.39 | 222290.91 | 336252.8 | 224172.1 |
| ALPQPQNVTSLLGC(cam)TH | HexNAc4Hex5Fuc/HexNAc4Hex5 | 1120.158 | 1168.844 | 3 | 1497.168 | 1570.197 | 32.22 | 358862.97 | 502478.6 | 261153.6 |
| ALPQPQNVTSLLGC(cam)TH | HexNAc5Hex6Fuc/HexNAc5Hex6 | 1241.869 | 1290.555 | 3 | 1679.734 | 1752.763 | 32.05 | 24787.56 | 21756.85 | 13979.42 |
| ALPQPQNVTSLLGC(cam)TH | HexNAc6Hex7Fuc/HexNAc6Hex7 | 1363.58 | 1412.266 | 3 | 1862.3 | 1935.329 | 31.95 | 9160.63 | 10837.99 | 6689.204 |
| DLQSLEDILHQVENK | HexNAc4Hex5Fuc/HexNAc4Hex5 | 1135.168 | 1183.854 | 3 | 1519.682 | 1592.711 | 19.76 | 46406.068 | 134156.3 | 58168.16 |
| DQC(cam)IVDDITYNVNDTFHK | HexNAc4Hex5Fuc/HexNAc4Hex5 | 1273.861 | 1322.547 | 3 | 1727.721 | 1800.75 | 32.68 | 15905.46 | 15056.4 | 13280.03 |
| DQC(cam)IVDDITYNVNDTFHK | HexNAc5Hex6Fuc/HexNAc5Hex6 | 1395.572 | 1444.258 | 3 | 1910.288 | 1983.317 | 32.67 | 16594.723 | 16875.31 | 12020.9 |
| EHEGAIYPDNTTDFQR | HexNAc4Hex5Fuc/HexNAc4Hex5 | 879.611 | 916.125 | 4 | 1050.768 | 1099.454 | 19.5 | 1099902 | 1400967 | 1181306 |
| EHEGAIYPDNTTDFQR | HexNAc5Hex6Fuc/HexNAc5Hex6 | 970.894 | 1007.409 | 4 | 1172.479 | 1221.165 | 19.31 | 36378.466 | 51039.93 | 43231.07 |
| EHEGAIYPDNTTDFQR | HexNAc6Hex7Fuc/HexNAc6Hex7 | 1062.177 | 1098.692 | 4 | 1294.19 | 1342.876 | 19.12 | 40618.116 | 27621.16 | 46102.86 |
| ELHHLQEQNVSNAFLDK | HexNAc4Hex5Fuc/HexNAc4Hex5 | 911.902 | 948.416 | 4 | 1093.823 | 1142.508 | 18.45 | 65313.619 | 191405.9 | 131598.3 |
| ELHHLQEQNVSNAFLDK | HexNAc5Hex6Fuc/HexNAc5Hex6 | 1003.185 | 1039.699 | 4 | 1215.533 | 1264.219 | 18.32 | 166818.05 | 500385 | 397290.4 |
| ELHHLQEQNVSNAFLDK | HexNAc6Hex7Fuc/HexNAc6Hex7 | 1094.4678 | 1130.982 | 4 | 1337.244 | 1385.93 | 18.21 | 6002.83 | 49231.36 | 42164.2 |
| FGC(cam)EIENNR | HexNAc5Hex6Fuc/HexNAc5Hex6 | 1042.741 | 1091.427 | 3 | 1381.042 | 1454.071 | 28.13 | 108708.81 | 40864.21 | 48401.32 |
| FNLTETSEAEIHQSFQHLLR | HexNAc5Hex6Fuc/HexNAc5Hex6 | 1097.733 | 1134.247 | 4 | 1341.6 | 1390.283 | 39.42 | 188343.21 | 909450.5 | 269866.7 |
| IC(cam)DLLVANNHFAHFFAPQNLTNMNK | HexNAc4Hex5Fuc/HexNAc4Hex5 | 911.207 | 940.418 | 5 | 1047.474 | 1083.988 | 30.21 | 111580.17 | 84752.74 | 101153.8 |
| ISEENETTC(cam)YMGK | HexNAc4Hex5Fuc/HexNAc4Hex5 | 1062.082 | 1110.768 | 3 | 1410.054 | 1483.083 | 16.83 | 108203.02 | 200884.1 | 286774.7 |
| LANLTQGEDQYYLR | HexNAc4Hex5Fuc/HexNAc4Hex5 | 1102.81 | 1151.496 | 3 | 1471.145 | 1544.174 | 26.37 | 140960.21 | 194473.3 | 170119.1 |
| LANLTQGEDQYYLR | HexNAc5Hex6Fuc/HexNAc5Hex6 | 1224.52 | 1273.207 | 3 | 1654.711 | 1726.74 | 26.11 | 129909.18 | 158968.8 | 109131.6 |
| LANLTQGEDQYYLR | HexNAc6Hex7Fuc/HexNAc6Hex7 | 1346.231 | 1394.917 | 3 | 1836.277 | 1909.306 | 25.96 | 10960.695 | 12733.24 | 12773.08 |
| LDAPTNLQFVNETDSTVLVR | HexNAc4Hex5Fuc/HexNAc4Hex5 | 964.438 | 1000.953 | 4 | 1163.871 | 1212.557 | 34.4 | 13116.089 | 24452.96 | 25186.49 |
| LGAC(cam)NDTLQQLMEVFK | HexNAc4Hex5Fuc/HexNAc4Hex5 | 1163.835 | 1212.521 | 3 | 1562.683 | 1635.711 | 45.74 | 96815.12 | 94736.97 | 104882.5 |
| LGAC(cam)NDTLQQLMEVFK | HexNAc5Hex6Fuc/HexNAc5Hex6 | 1285.546 | 1334.232 | 3 | 1745.249 | 1818.278 | 45.29 | 14348.926 | 23864.36 | 20572.07 |
| LGNWSAMPSC(cam)K | HexNAc4Hex5Fuc/HexNAc4Hex5 | 958.386 | 1007.072 | 3 | 1254.509 | 1327.538 | 22.59 | 84008.545 | 100469.1 | 38520.73 |
| LGNWSAMPSC(cam)K | HexNAc5Hex6Fuc/HexNAc5Hex6 | 1080.097 | 1128.782 | 3 | 1437.075 | 1510.104 | 22.59 | 38985.369 | 93065.39 | 21342.94 |
| LGNWSAMPSC(cam)K | HexNAc6Hex7Fuc/HexNAc6Hex7 | 1201.807 | 1250.493 | 3 | 1619.641 | 1692.67 | 22.35 | 11509.573 | 33250.57 | 16624.77 |
| LPTQNITFQTESSVAEQEAEFQSPK | HexNAc4Hex5Fuc/HexNAc4Hex5 | 1108.74 | 1145.253 | 4 | 1356.272 | 1404.958 | 32.07 | 94491.907 | 102663.1 | 34267.43 |
| LPTQNITFQTESSVAEQEAEFQSPK | HexNAc5Hex6Fuc/HexNAc5Hex6 | 1200.022 | 1236.536 | 4 | 1477.983 | 1526.669 | 31.87 | 77934.934 | 149639 | 97153.68 |
| LPTQNITFQTESSVAEQEAEFQSPK | HexNAc6Hex7Fuc/HexNAc6Hex7 | 1291.305 | 1327.82 | 4 | 1599.694 | 1648.38 | 31.72 | 21470.555 | 40034.63 | 65667.65 |
| LQAPLNYTEFQKPIC(cam)LPSK | HexNAc4Hex5Fuc/HexNAc4Hex5 | 968.197 | 1004.711 | 4 | 1168.882 | 1217.569 | 34.05 | 178349.59 | 44088.9 | 41627.57 |
| LQAPLNYTEFQKPIC(cam)LPSK | HexNAc5Hex6Fuc/HexNAc5Hex6 | 1059.48 | 1095.994 | 4 | 1290.593 | 1339.279 | 33.85 | 24745.708 | 22009.63 | 32077.07 |
| NLSMPLLPADFHK | HexNAc4Hex5Fuc/HexNAc4Hex5 | 1035.791 | 1084.477 | 3 | 1370.617 | 1443.646 | 34.35 | 369487.53 | 260973.1 | 91111.11 |
| QVFPGLNYC(cam)TSGAYSNASSTDSASYYPLTGDTR | HexNAc4Hex5Fuc/HexNAc4Hex5 | 1294.043 | 1330.558 | 4 | 1603.345 | 1652.031 | 35.04 | 4741.323 | 25888.34 | 11434.47 |
| SLTFNETYQDISELVYGAK | HexNAc4Hex5Fuc/HexNAc4Hex5 | 1267.552 | 1316.238 | 3 | 1718.258 | 1791.287 | 46.19 | 29534.888 | 69845.5 | 33932.53 |
| SWPAVGNC(cam)SSALR | HexNAc4Hex5Fuc/HexNAc4Hex5 | 1009.755 | 1058.441 | 3 | 1331.563 | 1404.591 | 24.22 | 4001414.8 | 2854133 | 2329995 |
| SWPAVGNC(cam)SSALR | HexNAc5Hex6Fuc/HexNAc5Hex6 | 1131.466 | 1180.151 | 3 | 1514.129 | 1587.158 | 23.92 | 581478.1 | 598990.2 | 358282.1 |
| SWPAVGNC(cam)SSALR | HexNAc6Hex7Fuc/HexNAc6Hex7 | 1253.1763 | 1301.862 | 3 | 1697.695 | 1769.724 | 23.78 | 50194.542 | 74730.98 | 44858.33 |
| VC(cam)QDC(cam)PLLAPLNDTR | HexNAc4Hex5Fuc/HexNAc4Hex5 | 1132.147 | 1180.833 | 3 | 1515.154 | 1588.18 | 29.71 | 56974.136 | 42961.52 | 40571.29 |
| VC(cam)QDC(cam)PLLAPLNDTR | HexNAc5Hex6Fuc/HexNAc5Hex6 | 1253.86 | 1302.54 | 3 | 1697.72 | 1770.75 | 29.39 | 146951.5 | 368503.7 | 157980.5 |
| VC(cam)QDC(cam)PLLAPLNDTR | HexNAc6Hex7Fuc/HexNAc6Hex7 | 1375.57 | 1424.25 | 3 | 1880.28 | 1953.31 | 29.38 | 16232.114 | 51846.21 | 25500.91 |
| VDKDLQSLEDILHQVENK | HexNAc4Hex5Fuc/HexNAc4Hex5 | 937.1752 | 973.69 | 4 | 1127.52 | 1176.206 | 27.08 | 79862.323 | 85159.43 | 75367.18 |
| VGQLQLSHNLSLVILVPQNLK | HexNAc4Hex5Fuc/HexNAc4Hex5 | 1312.653 | 1361.334 | 3 | 1785.911 | 1858.934 | 42.36 | 36021.829 | 33956.68 | 40458.39 |
| VSNQTLSLFFTVLQDVPVR | HexNAc4Hex5Fuc/HexNAc4Hex5 | 1262.592 | 1311.278 | 3 | 1710.818 | 1783.848 | 55.74 | 42311.645 | 76258.23 | 119544.2 |
| VSNQTLSLFFTVLQDVPVR | HexNAc6Hex7Fuc/HexNAc6Hex7 | 1384.303 | 1433 | 3 | 1893.385 | 1966.414 | 55.23 | 11499.932 | 18969.14 | 25957.01 |

| Intensity HCV | Intensity NASH | Intensity Healthy | Intensity ALD | Intensity HBV | Intensity HCV | Intensity NASH |
|---|---|---|---|---|---|---|
| | | Non-Fucosylated | | | | |
| 1258819 | 1794829 | 9562145.6 | 5295233.69 | 5711100.69 | 7485535.3 | 6899278.6 |
| 240186.8 | 348510.5 | 3370988.6 | 1667572.51 | 1751785.24 | 1637203.9 | 2014067.8 |
| 275638.5 | 570737.6 | 33070569 | 15871836.9 | 18340065.3 | 18797898 | 26123871 |
| 13263.82 | 25608.41 | 2930078.6 | 1225157.55 | 1403591.71 | 1805331.2 | 2207122.9 |
| 10635.3 | 12344.65 | 153371.84 | 54937.931 | 60716.734 | 83413.248 | 111397.11 |
| 66571.94 | 125458.2 | 1999004.9 | 948753.797 | 1426899.32 | 1791230.1 | 2148461 |
| 16464.83 | 26385.13 | 687523.28 | 241115.035 | 608744.447 | 304650.03 | 374592.08 |
| 17880.75 | 23073.29 | 114830.53 | 45174.845 | 96704.835 | 56284.587 | 57963.589 |
| 1234467 | 1856934 | 1421769.7 | 1003633.45 | 1187268.88 | 1287452.7 | 1426810.7 |
| 41169.8 | 76978.61 | 2209228.8 | 1255341.95 | 1771959.07 | 2630817 | 2209169 |
| 25268.22 | 29474.88 | 134099.47 | 77332.325 | 125214.453 | 346787.84 | 178496.66 |
| 151691.7 | 193211.5 | 8725942.1 | 5268837.77 | 5245144.9 | 6041449.7 | 7921395.1 |
| 430844.7 | 489665.5 | 853779.56 | 537256.849 | 510262.422 | 866483.04 | 964781.82 |
| 48557.26 | 47069.91 | 21052.512 | 33059.231 | 21952.016 | 50322.082 | 47929.001 |
| 53217.63 | 76527.39 | 244040.09 | 178855.345 | 187351.143 | 297882.55 | 254187.46 |
| 194958.5 | 389337.7 | 4248750.7 | 4381366.29 | 3264320.9 | 3007786.5 | 3617108.1 |
| 99968.81 | 95903.15 | 47748.935 | 143806.52 | 96545.725 | 202715.44 | 197320.15 |
| 70528.54 | 213901 | 1656825.6 | 1183029.24 | 1896553.45 | 759706.4 | 1266790.1 |
| 186142 | 208252 | 2815893.2 | 918270.957 | 1164575.71 | 1148867 | 1493843.1 |
| 128023.4 | 174219.2 | 2429536.3 | 595394.047 | 917207.611 | 1141123.4 | 1255543.2 |
| 12371.49 | 13707.82 | 253740.8 | 66203.773 | 107682.877 | 145593.84 | 162198.08 |
| 21238.63 | 35355.58 | 143423.95 | 83763.294 | 66548.746 | 79467.977 | 100230.63 |
| 98662.69 | 70565.69 | 6975275.9 | 4881451.37 | 7215600.2 | 6501519.7 | 5892678.2 |
| 21920.28 | 16437.44 | 231196.1 | 101441.503 | 246128.341 | 159223.37 | 239507.04 |
| 64417.8 | 174606.5 | 10160568 | 4637672.14 | 3611284.86 | 6666697.1 | 10322577 |
| 51947.79 | 87519.46 | 287061.63 | 116503.998 | 91136.966 | 181915.79 | 269229.61 |
| 33242.36 | 39242.1 | 47560.447 | 34279.912 | 27779.533 | 88789.665 | 110468.92 |
| 27580.73 | 29262.42 | 1174057.9 | 628902.656 | 725141.363 | 723153.73 | 891879.34 |
| 57406.42 | 87521.83 | 913128.26 | 608322.804 | 601748.47 | 897062.28 | 880887.4 |
| 33666.81 | 23237.4 | 61367.458 | 27607.489 | 17084.647 | 32839.86 | 33033.723 |
| 60372.72 | 137576.7 | 1784943.5 | 341085.024 | 608842.567 | 625578.63 | 824867.28 |
| 29112.24 | 33030.78 | 410233.24 | 145192.582 | 197712.613 | 335656.62 | 310574.21 |
| 206417.6 | 403239.3 | 5214607.4 | 1394983.95 | 1182128.47 | 2326775.2 | 3077507.9 |
| 22129.84 | 23838.07 | 633791.44 | 452501.306 | 310162.133 | 489378.94 | 559074.17 |
| 26628.29 | 43869.65 | 3388881.4 | 1523264.09 | 2190558.46 | 2091635.5 | 2259822.3 |
| 2213730 | 4125403 | 18725139 | 6896897.19 | 8872987.35 | 8429225.7 | 12180218 |
| 425041.4 | 743661.8 | 6862788.2 | 2616842.59 | 3264933.5 | 4192918.2 | 4893616.1 |
| 61539.21 | 106539 | 688750.39 | 237586.406 | 295193.329 | 524571.51 | 554298.2 |
| 30153.78 | 57640.33 | 949498.42 | 547592.224 | 593393.407 | 645891.74 | 826903 |
| 242224.7 | 327213.1 | 4123457.6 | 2328976.73 | 2372881.15 | 4019685.6 | 3397923.3 |
| 36754.27 | 45598.26 | 204441.71 | 118042.07 | 134992.822 | 231901.37 | 227352.47 |
| 163109.5 | 150468.3 | 370292.75 | 125191.236 | 168919.917 | 200881.43 | 229299.54 |
| 44536.04 | 39867.16 | 663713.38 | 307441.22 | 325468.039 | 468288.34 | 361190.32 |
| 152381.7 | 112594.5 | 133431.46 | 273346.893 | 417969.33 | 811238.25 | 594753.74 |
| 42820.29 | 36898.41 | 16054.501 | 11685.407 | 20261.734 | 60850.065 | 35414.806 |

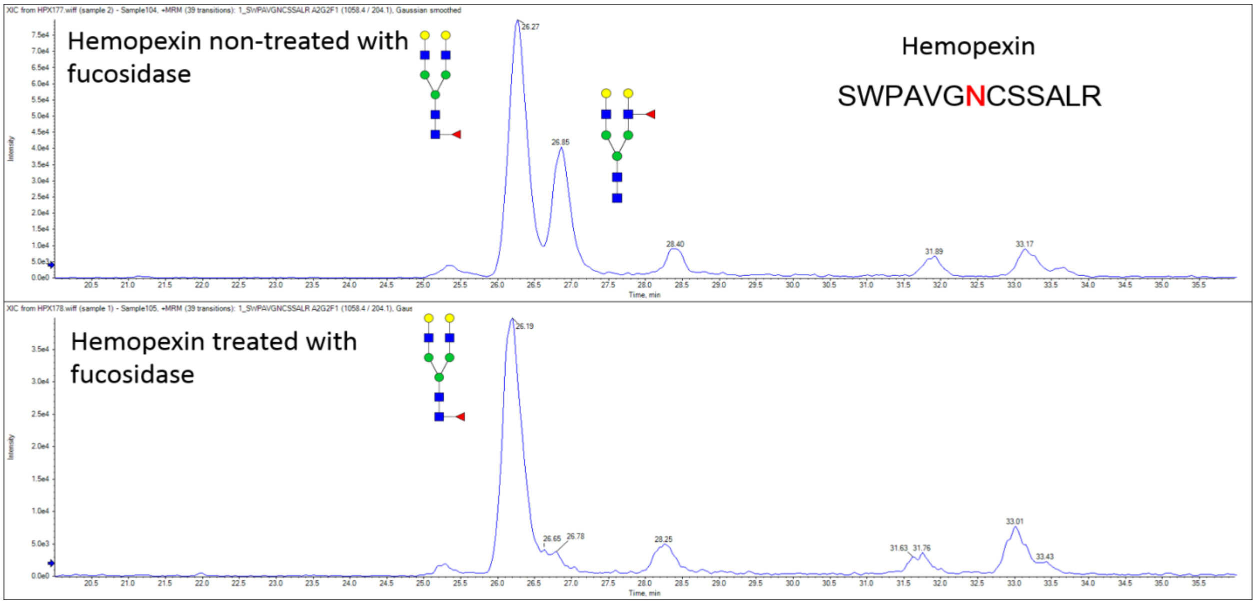


**Supplemental Figure 1:** HILIC chromatography of the glycopeptide of hemopexin treated with fucosidase confirms cleavage of the outer-arm fucose with > 98% efficiency.
